# Respirometry dataset for oxygen consumption measurements in carabid beetles

**DOI:** 10.64898/2026.04.28.720111

**Authors:** Sina Remmers, Kathrin H. Dausmann

## Abstract

**Overview:** This dataset originates from a preliminary respirometry study on carabid beetles from the Elbe Estuary (Northern Germany), encompassing species from freshwater and saltmarsh habitats along a salinity gradient. The study was designed to establish and validate a workflow for measuring oxygen consumption, including chamber setup, sensor recording, drift correction, and calculation of absolute and mass-specific metabolic rates. Oxygen consumption was measured for five species (*Carabus auratus, Carabus granulatus, Limodromus assimilis, Poecilus versicolor* and *Pterostichus niger*) using sealed glass vials connected to an optical oxygen system. The dataset provides individual-level measurements and serves primarily as a methodological reference for future respirometry studies on ground-dwelling arthropods. The O_2_ consumption rates of carabid beetles showed interspecific differences and followed metabolic scaling theory, revealing an inverse relationship between body mass and mass-specific metabolic rates across species (Figure 3).

## Background

Oxygen consumption is widely used as a proxy for metabolic rate in ecophysiological studies of arthropods, because it provides a standardised measure of energy expenditure under controlled conditions and allows comparison across taxa, body sizes and environmental contexts (Nespolo et al., 2003). However, methodological challenges such as activity effects and measurement technique differences can complicate direct interpretation (Van Voorhies et al., 2008). In carabid beetles, these measurements are valuable for understanding physiological adaptation to habitat conditions, including temperature, moisture, and disturbance regimes (Alford et al., 2023). This is especially relevant for species from the Elbe estuarine marshlands, where tidal influence, periodic inundation and spatial variation in moisture create a dynamic heterogenous environment that may shape metabolic rates and physiological constraints (Butzek, 2014; Remmers & Dausmann, 2025). This pilot study focused on method development rather than test specific hypotheses. In particular, the study addressed several challenges of O_2_ consumption easurements, such as sensor drift, handling effects immediately after transfer to chambers, changing oxygen trajectories caused by intermittent activity and the need to account for beetle body volume when converting oxygen decline to consumed gas volume. Carabid beetles are known to show distinct respiratory patterns, including discontinuous gas exchange and spiracle regulation, which can intermittently reduce or pause gas exchange and produce non-linear oxygen traces if entire recording periods are analysed (Duncan & Dickman, 2001; Socha et al., 2008)

### Study design

Fieldwork was conducted at two capture sites along the Elbe estuary in northern Germany, representing freshwater and saltmarsh habitat conditions along the estuarine salinity gradient. Carabid beetles were sampled using live pitfall traps, following standard methods for ground-dwelling arthropods (Barber, 1931). Traps remained open for up to five days and contained paper towel soaked in red wine as an attractant, along with leaf litter and bark that provided shelter and acted as a flood buffer during heavy rain or inundation. Beetles larger than 5 mm were identified, and the most abundant species were retained for respirometry; all others were released at the capture site. Beetles were weighed individually before measurement and placed into separate sealed 20 mL glass vials equipped with integrated optical oxygen sensors connected to a FireStingO_2_ system (PyroScience GmbH, Aachen, Germany). Oxygen concentration and ambient temperature were recorded continuously for around 12 hours for each individual. Each measurement round included three sample vials with beetles and one empty control vial. The control was used to quantify oxygen drift not caused by beetle respiration and to correct minute-wise oxygen values of the sample vial accordingly. Measurements were scheduled to coincide with the animals’ inactive phase in order to reduce activity-related noise. Diurnal species, *C. auratus* and *P. versicolor*, were measured under dark conditions during the night, whereas nocturnal species, *C. granulatus, L. assimilis*, and *P. niger*, were measured under light conditions during the day (Klaiber et al., 2017) (Figure 2). For the conversion from oxygen concentration change in percentage to consumed oxygen volume in mL, species-specific average body volume was determined independently by water displacement using five individuals for each measured species from previous monitoring pitfall trap samples. Mean body volume for each species was then subtracted from the vial volume for subsequent calculations.

**Figure 1:**
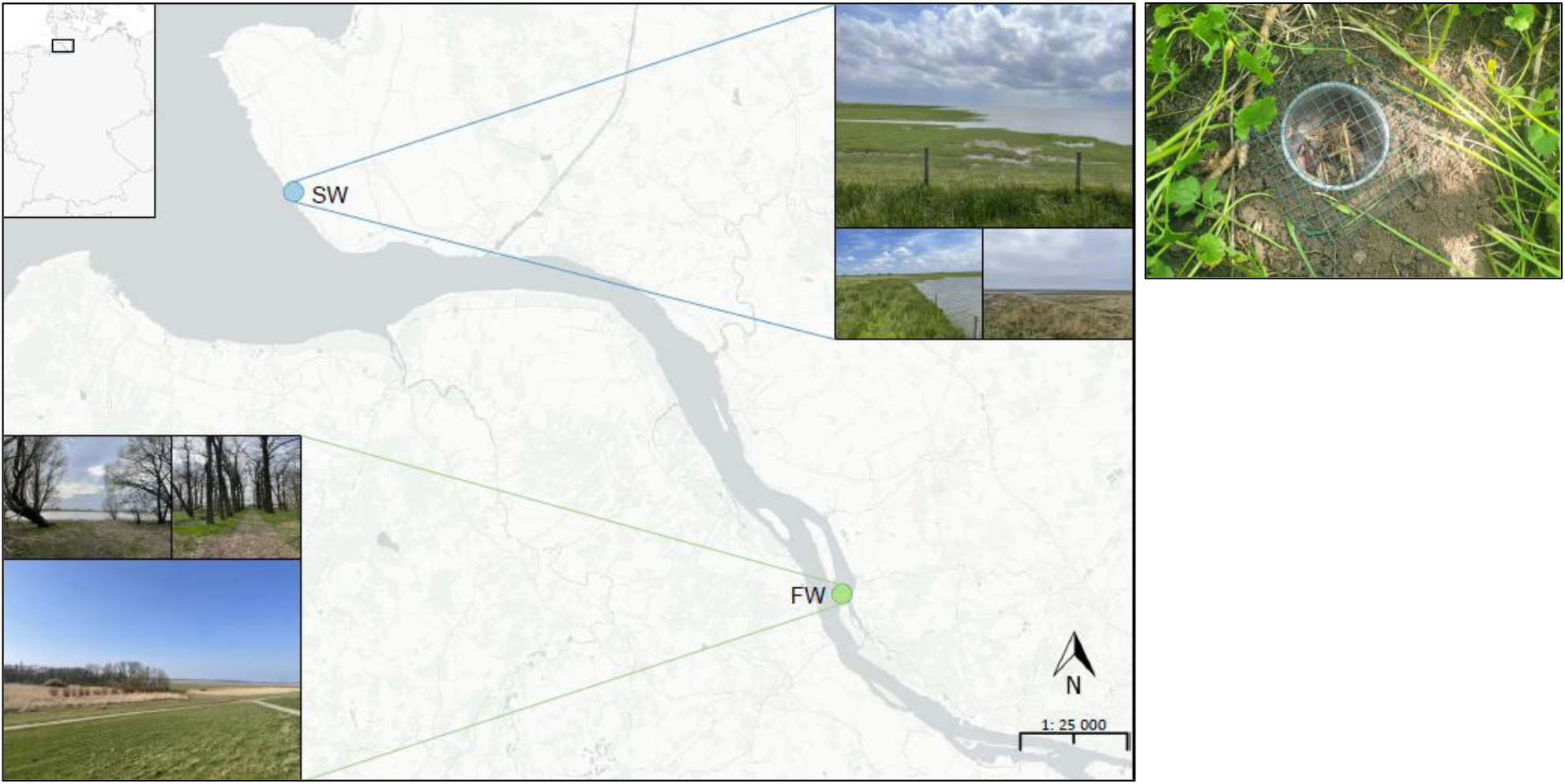
Map showing the locations of the capture sites in the freshwater marsh (FW; green) and the saltwater marsh (SW; blue). Photographs depict the characteristic habitats of each marsh type and a typical pitfall trap setup.

**Figure 2:**
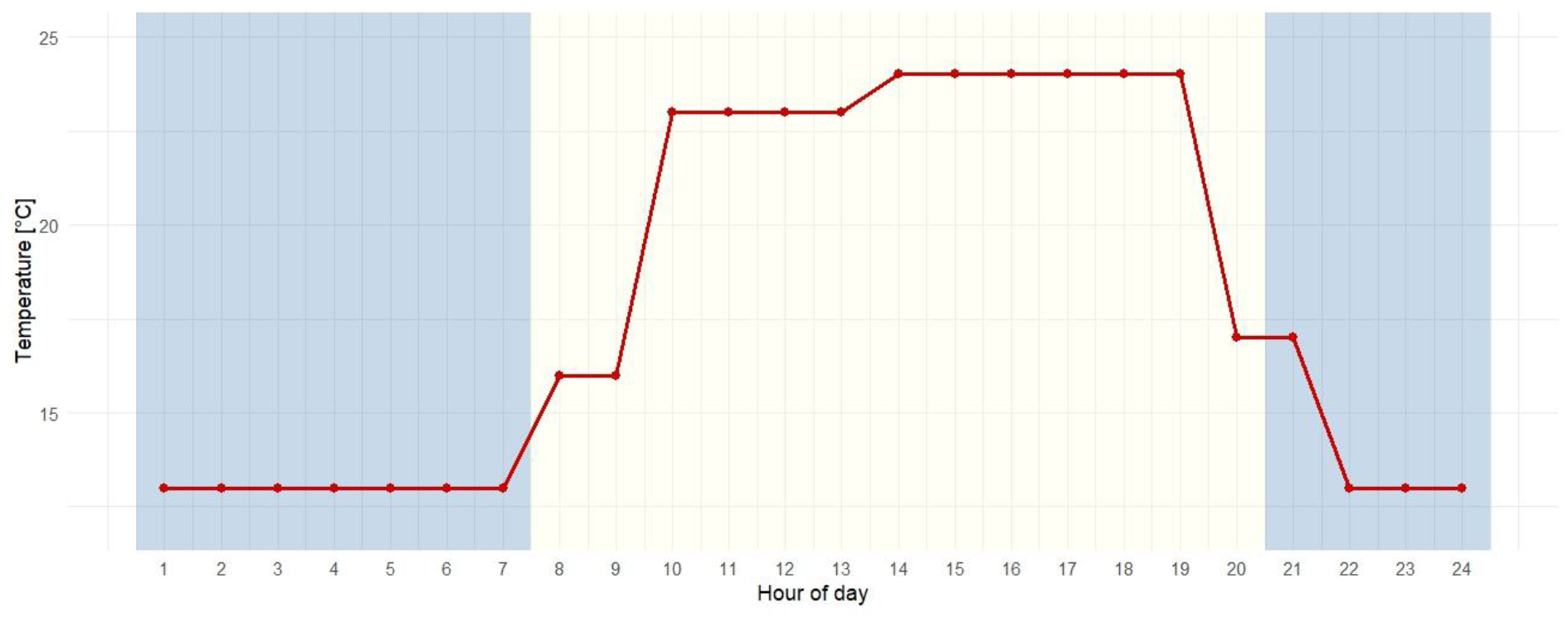
Simulated temperature cycle and day-night rhythm in the climate chamber. The temperature profile reflects average diurnal conditions of August, while the day–night rhythm was simulated by alternating the chamber lighting to reflect a natural photoperiod.

**Figure 3:**
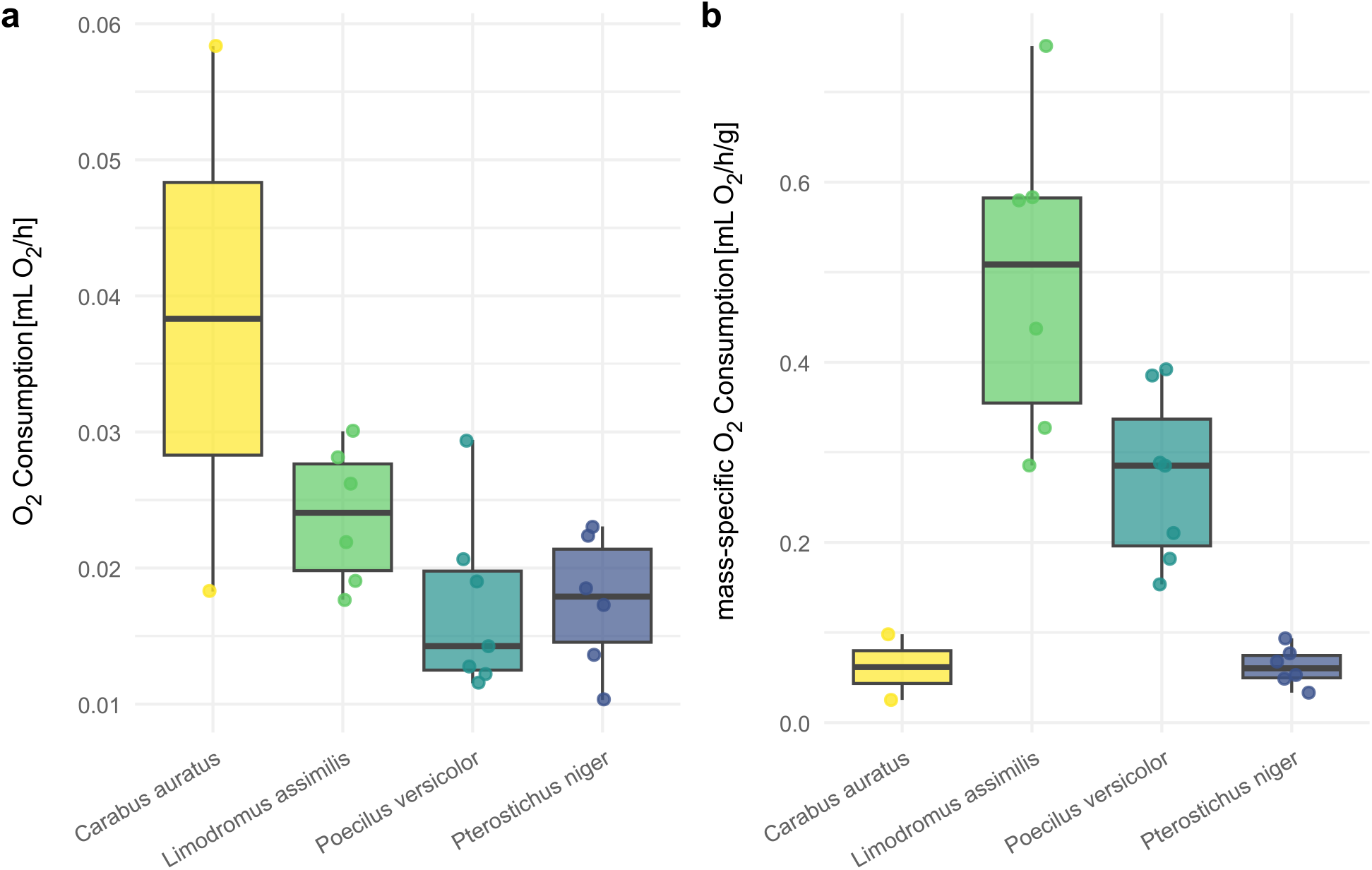
Oxygen consumption rates of Carabid species per (**a**) animal in [ml O_2_ h^−1^] and as (**b**) mass-specific consumption rate [ml O_2_ h^−1^ g^−1^]. Points represent mean oxygen consumption per individual (C. auratus: n = 2; L. assimilis: n = 6; P. versicolor: n = 7; P. niger: n = 6).

### Data processing

Raw oxygen data were aggregated to minute-wise mean oxygen concentrations for each individual. These values were corrected using the empty control vial from the same measurement round to remove background drift attributed to the sensor or external influences such as temperature fluctuation. We calculated the drift as the continuous relative change in the control compared to the baseline %O_2_ concentration at the start of each measurement, rather than subtracting absolute values, to avoid bias from small differences in baseline %O_2_ among channels. This drift correction was then applied individually to each specimen within the respective run. The first and last 30 minutes of each recording were excluded before further analysis. This step was intended to minimise artefacts caused by handling, chamber closure and external disturbance during or at the end of the run. After processing, linear periods with a stable decline in oxygen concentration were identified manually for each beetle. Because carabids can intermittently restrict gas exchange via spiracle regulation, producing non-linear O_2_ traces (Socha et al., 2008), only manually selected linear segments showing a steady decline in oxygen were analysed. Oxygen consumption was calculated for the selected interval as mL O_2_min^−1^, converted to mL O_2_ h^−1^, and standardised by body mass to obtain mL O_2_ h^−1g-1^. Mean body mass per beetle was defined as the average of pre- and post-measurement body mass.

### Dataset contents

The data should be interpreted primarily as a methodological reference. Sample sizes were limited and uneven across species, and the pilot study was not designed to support strong biological conclusions about interspecific metabolic differences. The dataset contains individual ID’s, species identity, timestamps from the selected measurement intervals, oxygen consumption rate per minute and per hour, mean body mass, and mass-specific oxygen consumption rate (Table 1). The dataset includes measurements for 22 individuals across five species: *Carabus auratus* (n = 2), *Carabus granulatus* (n = 1), *Limodromus assimilis* (n = 6), *Poecilus versicolor* (n = 7), and *Pterostichus niger* (n = 6). Across the 22 individuals measured, consumption rates generally showed interspecific differences (Figure 3). Hourly oxygen consumption rates ranged from 0.0103 to 0.0584 mL O_2_ h^−1^ across individuals, with highest absolute values observed in larger carabids (Figure 3a). Mass-specific oxygen consumption rates ranged from 0.0252 to 0.7512 mL O_2_ h^−1^ g^−1^, with the highest values in small species like *L. assimilis* and the lowest in larger ones such as *C. auratus* and *P. niger* (Figure 3b). These findings are consistent with metabolic scaling theory, which predicts that larger carabids exhibit lower mass-specific metabolic rates than smaller species, as seen in the negative relationship between body mass and standardised oxygen consumption across the dataset (Gudowska et al., 2017).

**Table 1:** Individual oxygen consumption rates and body mass of six carabid beetle species measured in respirometry trials. Consumption rates are expressed per minute (mL O_2_ min^−1^), per hour (mL O_2_ h^−1^), and mass-specific per hour (mL O_2_ h^−1^ g^−1^). TS start and TS end mark the timestamps from the analysed time period of the measurement.

| ID | species | TS start | TS end | consumption<br>rate [per<br>min] | consumption<br>rate [per<br>hour] | mean<br>BM<br>[mg] | consumption<br>rate<br>[hour_g] |
| --- | --- | --- | --- | --- | --- | --- | --- |
| Car_012 | *C. auratus* | 04.11.24<br>20:08 | 04.11.24<br>21:41 | 0.0016 | 0.0183 | 725 | 0.0252 |
| Car_014 | *C. auratus* | 31.10.24<br>21:12 | 31.10.24<br>23:50 | 0.0051 | 0.0584 | 595 | 0.0981 |
| Car_117 | *C. granulatus* | 31.10.24<br>21:21 | 01.11.24<br>04:05 | 0.0015 | 0.0181 | 310 | 0.0583 |
| Lim_301 | *L. assimilis* | 30.10.24<br>12:35 | 30.10.24<br>13:37 | 0.0024 | 0.0281 | 48.5 | 0.5798 |
| Lim_302 | *L. assimilis* | 01.11.24<br>14:52 | 01.11.24<br>15:27 | 0.0025 | 0.03 | 40 | 0.7512 |
| Lim_303 | *L. assimilis* | 30.10.24<br>12:47 | 30.10.24<br>13:41 | 0.0016 | 0.0191 | 67 | 0.2854 |
| Lim_306 | *L. assimilis* | 04.11.24<br>14:59 | 04.11.24<br>16:35 | 0.0022 | 0.0263 | 45 | 0.5834 |
| Lim_309 | *L. assimilis* | 01.11.24<br>13:56 | 01.11.24<br>15:02 | 0.0018 | 0.0219 | 50 | 0.4374 |
| Lim_310 | *L. assimilis* | 30.10.24<br>09:00 | 30.10.24<br>10:00 | 0.0015 | 0.0177 | 54 | 0.3272 |
| Poe_035.1 | *P. versicolor* | 29.10.24<br>19:43 | 29.10.24<br>21:26 | 0.0017 | 0.0206 | 53.5 | 0.3853 |
| Poe_046 | *P. versicolor* | 28.10.24<br>19:44 | 28.10.24<br>22:54 | 0.0025 | 0.0294 | 75 | 0.3922 |
| Poe_065 | *P. versicolor* | 28.10.24<br>20:07 | 29.10.24<br>03:27 | 0.001 | 0.0123 | 80 | 0.1535 |
| Poe_313.2 | *P. versicolor* | 30.10.24<br>21:12 | 31.10.24<br>00:27 | 0.0016 | 0.0189 | 90 | 0.2105 |
| Poe_314 | *P. versicolor* | 28.10.24<br>20:59 | 29.10.24<br>01:45 | 0.0011 | 0.0127 | 70 | 0.1818 |
| Poe_315.2 | *P. versicolor* | 30.10.24<br>21:14 | 31.10.24<br>00:47 | 0.0012 | 0.0143 | 50 | 0.2853 |
| Poe_316 | *P. versicolor* | 30.10.24<br>21:38 | 31.10.24<br>04:37 | 0.001 | 0.0115 | 40 | 0.2884 |
| Pte_017.2 | *P. niger* | 04.11.24<br>17:18 | 04.11.24<br>18:21 | 0.0009 | 0.0103 | 310 | 0.0333 |
| Pte_079 | *P. niger* | 31.10.24<br>13:38 | 31.10.24<br>15:31 | 0.0019 | 0.0223 | 330 | 0.0677 |
| Pte_318 | *P. niger* | 29.10.24<br>11:48 | 29.10.24<br>12:26 | 0.0012 | 0.0136 | 280 | 0.0487 |
| Pte_326.2 | *P. niger* | 04.11.24<br>12:22 | 04.11.24<br>15:19 | 0.0015 | 0.0173 | 185 | 0.0935 |
| Pte_327 | *P. niger* | 31.10.24<br>12:47 | 31.10.24<br>14:10 | 0.0019 | 0.0231 | 300 | 0.0769 |
| Pte_329 | *P. niger* | 29.10.24<br>12:13 | 29.10.24<br>13:21 | 0.0016 | 0.0185 | 350 | 0.0529 |

### Reuse value

The dataset is being made publicly available because published data on oxygen consumption rates in carabid beetles remain comparatively scarce, despite their value for comparative physiology, ecological inference and methodological standardisation. Making these pilot measurements accessible helps preserve otherwise underrepresented data and may support future studies and broader comparative analyses of carabid respiratory physiology. The protocol described here directly provided the basis for a later temperature-based study in which carabid oxygen consumption was measured under controlled thermal treatments using an expanded and partly modified design. Because the paper is intended as a data-focused publication, emphasis should remain on study design, measurement chamber setup, recording conditions and processing decisions rather than on ecological interpretation of species differences. This makes the dataset citable as a methodological resource for future respirometry studies on beetles and other terrestrial arthropods.

